# Generation of “OP7 chimera” defective interfering particle preparations free of infectious influenza A virus that shows antiviral efficacy in mice

**DOI:** 10.1101/2023.08.16.553516

**Authors:** Tanya Dogra, Lars Pelz, Julia D. Boehme, Jan Küchler, Olivia Kershaw, Pavel Marichal-Gallardo, Maike Bälkner, Marc D. Hein, Achim D. Gruber, Dirk Benndorf, Yvonne Genzel, Dunja Bruder, Sascha Y. Kupke, Udo Reichl

## Abstract

Influenza A virus (IAV) defective interfering particles (DIPs) are considered as new promising antiviral agents. Conventional DIPs (cDIPs) contain a deletion in the genome and can only replicate upon co-infection with infectious standard virus (STV), during which they suppress STV replication. We previously discovered a new type of IAV DIP “OP7” that entails genomic point mutations and displays higher antiviral efficacy than cDIPs. To avoid safety concerns for the medical use of OP7 preparations, we developed a production system that does not depend on infectious IAV. We reconstituted a mixture of DIPs consisting of cDIPs and OP7 chimera DIPs, in which both harbor a deletion in their genome. To complement the defect, the deleted viral protein is expressed by the suspension cell line used for production in shake flasks. Here, DIP preparations harvested are not contaminated with infectious virions, and the fraction of OP7 chimera DIPs depended on the multiplicity of infection. Intranasal administration of OP7 chimera DIP material was well tolerated. A rescue from an otherwise lethal IAV infection and no signs of disease upon OP7 chimera DIP co-infection demonstrated the remarkable antiviral efficacy. The clinical development of this new class of broad-spectrum antiviral may contribute to pandemic preparedness.

## Introduction

Influenza A virus (IAV) is a major human pathogen. Infections cause annual epidemics, which lead to excessive morbidity and mortality [1]. When novel strains emerge, IAV infections may result in a severe pandemic, which is considered as an imminent threat. For instance, more than 40 million deaths were reported during the “Spanish flu” in 1918 [2]. Annual prophylactic vaccination is the most effective measure to prevent seasonal influenza infection [3]. Yet, the selection of strains as well as the manufacturing and release of a seasonal vaccines requires several months. Thus, small molecule drug antivirals are also used, for instance, to treat acute infections [1]. However, circulating human IAV strains have acquired resistance against many current antivirals [3]. Therefore, new broadly-acting antiviral treatment options should be considered not only to complement annual vaccination schemes but also to act as a first line of defense for pandemic preparedness.

Defective interfering particles (DIPs) are regarded as a promising new class of antivirals [4–17]. In particular, IAV-derived DIPs resulted in a high tolerability and antiviral efficacy in animal studies [5, 16, 18–25], and were therefore proposed as prophylactic and therapeutic antivirals [16, 25–27]. IAV DIPs typically contain a large internal deletion in one of the eight genomic viral RNA (vRNA) segments [4, 10, 16, 22, 25, 26, 28–30]. The missing genomic information results in the expression of a truncated viral protein [31]. Therefore, DIPs are defective in virus replication and cannot propagate in mammalian cells. In a co-infection with an infectious standard virus (STV), however, the missing gene function (i.e., the full-length (FL) protein) is provided, and DIPs are able to propagate. Interestingly, this results in a strong interference with STV replication. With respect to this antiviral effect, it is suggested that the short defective interfering (DI) vRNAs replicate faster and accumulate to higher levels than the FL vRNAs. Thereby, cellular and viral resources are depleted, which suppresses infectious virus replication [32–34]. DIP co-infections also result in a strong induction of the interferon (IFN) system [35–38], and it was shown that this stimulation of the innate immunity also contributes to their antiviral effect [25, 26, 35, 37]. As a consequence, IAV DIPs display a broad-spectrum antiviral activity that is not only directed against a wide range of IAV strains [16, 21, 25, 39, 40], but even against unrelated viruses, including SARS-CoV-2 [37, 38, 41].

Previously, we developed a cell culture-based production process [18, 42] for a well-known DIP called “DI244” that harbors a deletion in segment 1 (Seg 1) [25, 26]. DI244 is unable to express the viral polymerase basic protein 2 (PB2, encoded from Seg 1) and can be propagated in genetically engineered PB2-expressing Madin-Darby canine kidney (MDCK) suspension (MDCK-PB2(sus)) cells [18, 42, 43]. In addition, using a modified reverse genetics workflow for IAV that is specific for DIP rescue, purely clonal DI244 without STVs could be reconstituted for production [43]. Therefore, considering the use of DIPs as an antiviral, the absence of infectious STVs is expected to alleviate potential safety and regulatory concerns.

We previously discovered a new type of IAV DIP, called “OP7” that contains multiple point substitutions on segment 7 (Seg 7) vRNA instead of a large internal deletion [39]. OP7 showed a higher antiviral activity compared to Seg 1 conventional DIPs (cDIPs) including DI244 as shown in *in vitro* and *in vivo* experiments [18, 19, 37]. As the source of the defect in virus replication of OP7 is yet unknown, designing a cell line that could complement the defect of OP7 was not feasible, so far. Instead, we recently established a cell culture-based production process for OP7 in the presence of infectious STVs to complement the unknown defect [19]. However, infectious STVs had to be UV-inactivated, which also reduced antiviral activity of OP7. Moreover, even after UV treatment, the risk of a contamination with residual STVs should raise safety concerns with respect to medical application.

In the present study, we devised a genetically engineered cell culture-based production system for OP7, which does not require the addition of any infectious “STV”. Animal trials in mice suggest that the produced OP7 preparations can be used as a safe and potent antiviral, and further steps towards clinical development seems promising.

## Results

### Reconstitution of OP7 chimera DIPs without infectious STVs

Previously, OP7 was produced in cell culture in the presence of infectious STVs [19]. To obtain an OP7 seed virus without infectious STVs (STV-free), we modified a plasmid-based reverse genetics system for the reconstitution of Seg 1-derived cDIPs based on the IAV strain A/PR/8/34 (PR8) as described previously [43]. As Seg 1 cDIPs contain a large internal deletion in Seg 1 and are unable to express the viral PB2 protein, the STV-free reconstitution of purely clonal Seg 1 cDIPs requires PB2-expressing cells (Fig. 1A, B and C). Here, a co-culture of adherent PB2-expressing human embryonic kidney (HEK-293T-PB2(adh)) and MDCK-PB2(adh) cells (Fig. 1B) were co-transfected with eight plasmids encoding for the deleted Seg 1 and the remaining seven wild-type (WT) segments (Fig. 1A). After reconstitution, such Seg 1 cDIPs (Fig. 1C) could be propagated in cell culture using PB2-expressing cells, as shown previously [18, 42, 44].

**Figure 1:**
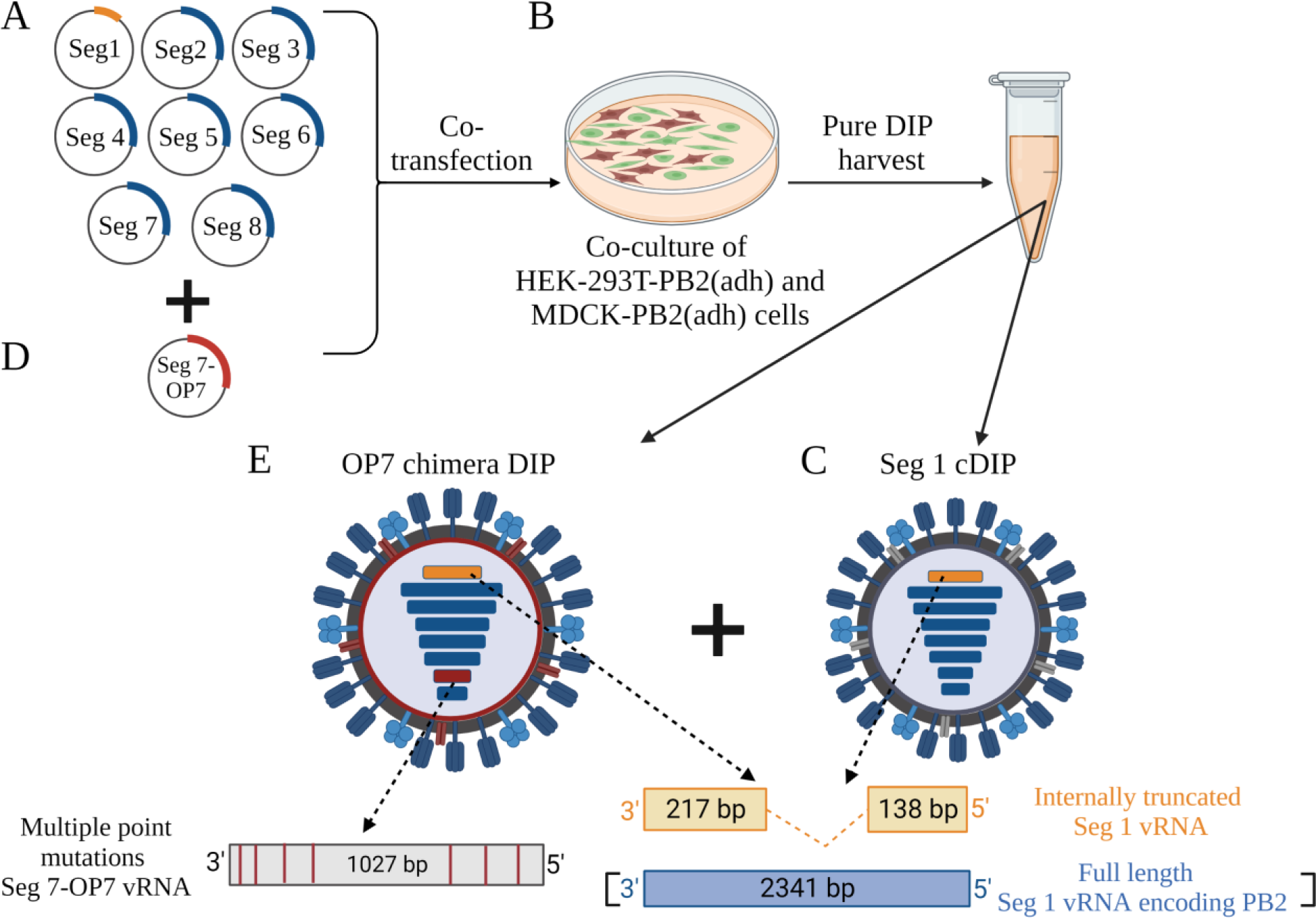
Plasmid-based reconstitution of OP7 chimera DIPs free of infectious STVs. Rescue of Seg 1 cDIPs. The reverse genetics system comprises (**A**) eight plasmids that encode for a deleted Seg 1 vRNA and Seg 2-8 WT vRNAs. (**B)** Co-transfection of a co-culture of PB2-expressing HEK-293T-PB2(adh) (high transfection efficiency) and MDCK-PB2(adh) (high virus titers) cells results in reconstitution of (**C**) purely clonal Seg 1 cDIP free of infectious STVs. Rescue of OP7 chimera DIPs: addition of a (**D**) ninth plasmid that encodes for the mutated Seg 7-OP7 vRNA results in the reconstitution of a mixture of DIPs including (**E**) OP7 chimera DIPs and (**C**) Seg 1 cDIPs. This mixture of viruses can be propagated in MDCK-PB2(sus) cells (Fig. 2). Image was created with BioRender.com.

In order to reconstitute OP7 chimera DIPs without infectious STVs, we added a ninth plasmid encoding for the mutated Seg 7 of OP7 (Seg 7-OP7) (Fig. 1D) for transfection. This resulted in the rescue of a population of two types of DIPs: (i) Seg 1 cDIPs (Fig. 1C) and (ii) OP7 chimera DIPs (Fig. 1E). Accordingly, the Seg 1 cDIPs contained a truncated Seg 1 vRNA and seven WT vRNAs (Fig. 1C) and OP7 chimera DIPs contained Seg 7-OP7 vRNA, a truncated Seg 1 vRNA, and the remaining six WT vRNAs (Fig. 1E). Owing to a deletion in Seg 1 (encoding for PB2), both DIPs could be propagated in MDCK-PB2(sus) cells (Fig. 2). Furthermore, it has to be assumed that the OP7 chimera DIPs (Fig. 1E) are defective in virus replication in PB2-expressing cells, as they contain the mutated and defective Seg 7-OP7 vRNA. Accordingly, for propagation, OP7 chimera DIPs require complementation with Seg 1 cDIPs (Fig. 1C) as they provide the functional Seg 7-WT vRNA (Seg 7-WT). Moreover, as both DIPs (Fig. 1C and E) are replication deficient in non-PB2 expressing cells, we eliminate the need for post-production UV inactivation due to the lack of infectious STVs. Note that the deleted Seg 1 sequence used in the present study was previously identified by us (“Seg 1 gain”), where corresponding DIPs showed a superior *in vitro* interfering efficacy compared to the well-known DI244 [44]. The absence of infectious STVs in the produced OP7 chimera DIP material was evaluated by two serial infection passages in adherent WT MDCK (MDCK(adh)) cells. Both passages showed no virus titer, thus confirming no infectious STV replication (innocuity assay, data not shown).

**Figure 2:**
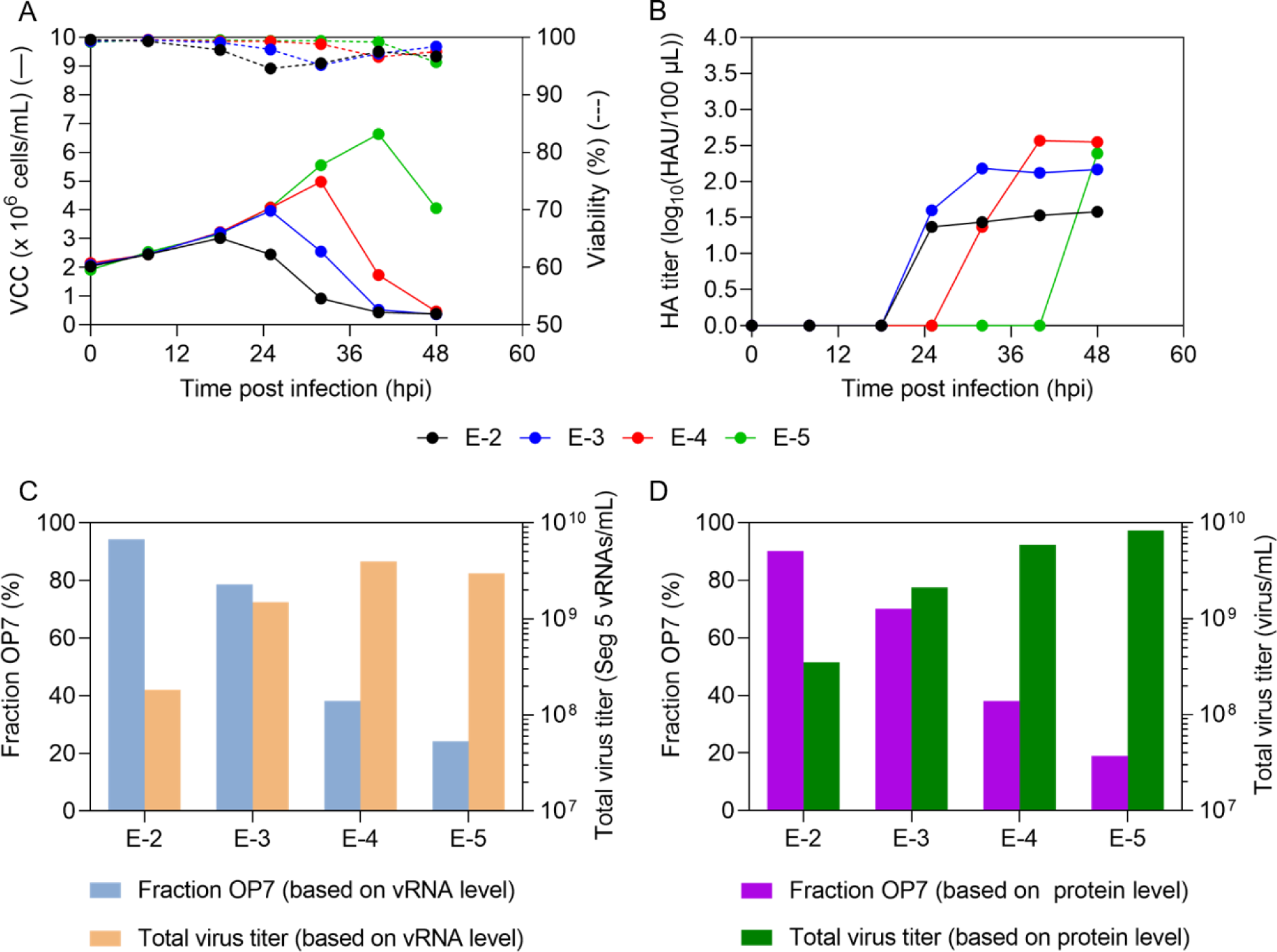
Cell culture-based production of OP7 chimera DIPs in shake flasks. Genetically engineered MDCK-PB2(sus) cells cultivated in 125 mL shake flasks (50 mL working volume), were infected at MOIs ranging from 1E-2 to 1E-5 after a complete medium exchange. (**A**) VCC and viability. (**B**) HA titer. (**C**) Fraction of OP7 chimera DIPs (calculated based on extracellular Seg 7-OP7 and Seg 7-WT vRNA concentrations, quantified by RT-qPCR). Total virus concentration is indicated by the extracellular Seg 5 vRNA concentration. (**D**) Fraction of OP7 chimera DIPs (calculated based on extracellular M1-OP7 and M1-WT viral protein concentrations, quantified by MS). Total virus concentration was calculated from the protein concentration of NP, encoded by Seg 5. The optimal harvest time points (MOI 1E-2: 25 hpi, 1E-3: 32 hpi, 1E-4: 40 hpi, 1E-5: 48 hpi) were analyzed for C and D. The figure depicts the results of one experiment.

In summary, we reconstituted OP7 chimera DIPs in a mixture with Seg 1 cDIPs without addition of any infectious STVs. Seg 1 cDIPs can complement the defect of the OP7 chimera DIPs in PB2-expressing MDCK cells, allowing for cell culture-based production.

### Cell culture-based production of OP7 chimera DIPs in shake flasks shows strong dependence on the multiplicity of infection

As indicated above, OP7 chimera DIPs are defective in virus replication in MDCK-PB2(sus) cells and propagation requires co-infection with Seg 1 cDIPs. Previously, in a similar production system, we produced OP7 in the presence of infectious STVs in WT MDCK(sus) cells. As expected for virus production, total virus yields were multiplicity of infection (MOI) dependent [19]. High MOI conditions increased the likelihood of co-infection of STVs and OP7, resulting in preferential production of OP7 that suppressed STV propagation and thus, total virus yields. In contrast, in a low MOI scenario, more single hit infections occurred. Therefore, STV growth occurred predominantly and the propagation-incompetent OP7 was out-diluted, resulting in higher total virus titers with a concomitant lower fraction of OP7 [19]. We expected that our new DIP mixture containing OP7 chimera DIPs and Seg 1 cDIPs shows the same MOI dependency in PB2-expressing cells. Thus, to optimize total virus yields and the fraction of OP7 chimera DIPs within the mixture, we performed infections in shake flasks using MDCK-PB2(sus) cells at different MOIs ranging from 1E-2 to 1E-5 (Fig. 2).

After infection at 2.1 × 10^6^ cells/mL, cells continued to grow (Fig. 2A). The viable cell concentration (VCC) post infection at a MOI of 1E-2 peaked fastest (3.0 × 10^6^ cells/mL, 18 hpi) before virus-induced cell lysis became evident by a decrease in VCC. With decreasing MOIs, the maximum VCC increased and cell death started later. As expected, the hemagglutinin (HA) titer (indicating total virus yield) reached lower values at higher MOIs relative to lower MOIs (Fig. 2B), likely due to the inhibition caused by increasing accumulation of OP7 chimera DIPs towards higher MOIs. This is in line with greater fractions of OP7 chimera DIPs at higher MOIs (Fig. 2C and D). For instance, we found a fraction of OP7 chimera DIPs of 94.4% (MOI 1E-2) and 24.2% (MOI 1E-5), calculated based on the extracellular vRNA concentration of Seg 7-OP7 and Seg 7 of the WT virus quantified by RT-qPCR (Fig. 2C). In addition, quantification of IAV proteins by a new mass spectrometry (MS) method developed in our group [45] showed similar fractions of 90.2% (MOI 1E-2) and 19.0% (MOI 1E-5). Here the fraction was calculated based on the concentration of the extracellular matrix protein 1 (M1, encoded on Seg 7) of OP7 (M1-OP7) and of the WT virus (M1-WT) (Fig. 2D). Further, the total virus concentration (as indicated by the extracellular Seg 5 vRNA concentration (Fig. 2C) and calculated from the NP concentration (Fig. 2D)) showed higher values for lower MOIs in line with HA titers (Fig. 2B). Previously, biological activity of IAV particles decreased over time as seen by a drop in the infectious virus titers towards late process times [46], which is important for selecting the optimal harvest time point (Fig 2B). Accordingly, for DIP harvesting, we selected the time point at which the HA titer almost plateaued to ensure maximum virus release and biological activity of the DIPs (MOI 1E-2: 25 hpi, MOI 1E-3: 32 hpi, MOI 1E-4: 40 hpi, MOI 1E-5: 48 hpi). Furthermore, harvesting was performed no later than the onset of cell death (Fig. 2A) to avoid excessive levels of cell debris and host cell DNA in the supernatant that would otherwise interfere with the subsequent downstream purification process. Further, no apparent accumulation of other DI vRNAs harboring a deletion in Seg 2-8 was detected by PCR (Fig. 3).

**Figure 3:**
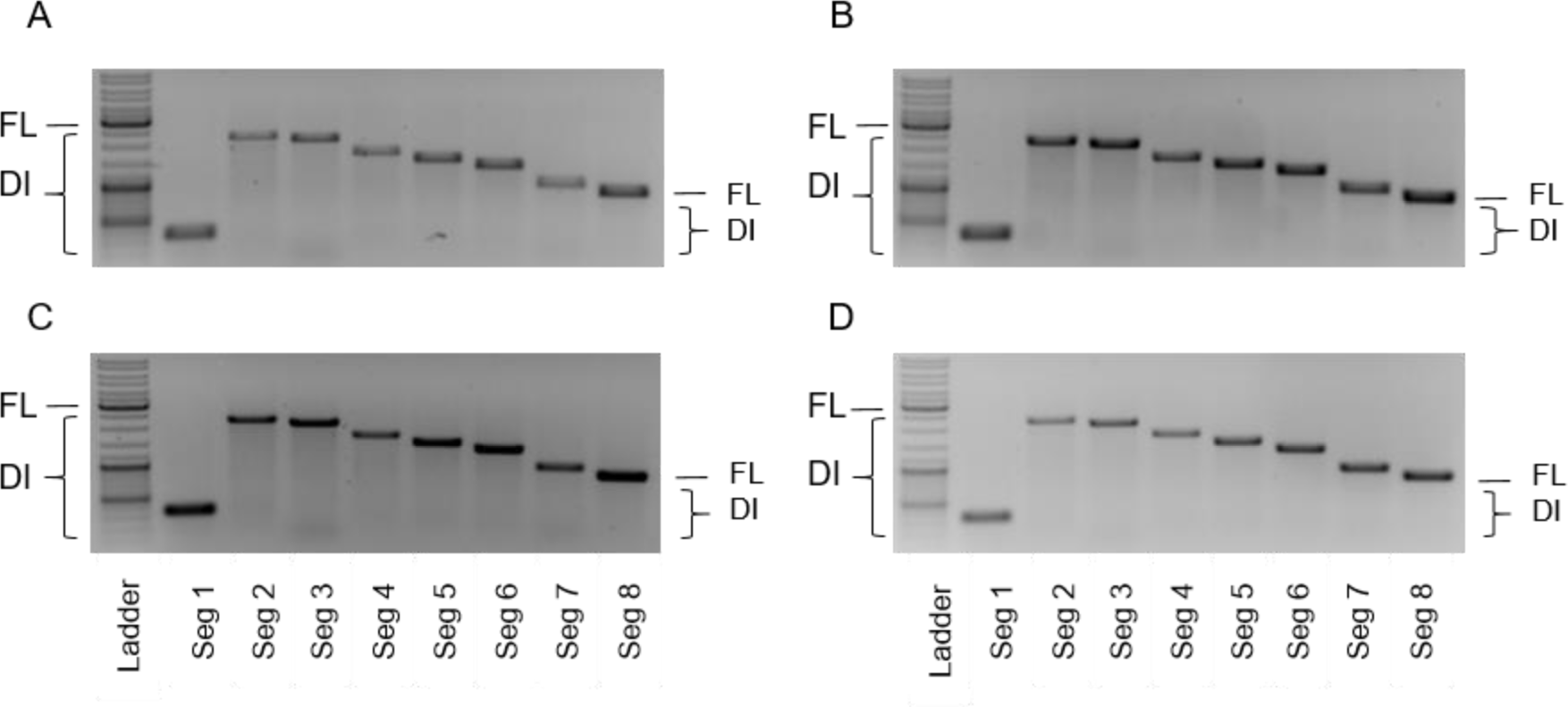
Purity of produced OP7 chimera DIP material with respect to contaminating DIPs. OP7 chimera DIPs were produced at different MOIs in shake flasks (Fig. 2). Samples from 48 hpi were subjected to segment-specific RT-PCR and gel electrophoresis. (**A**) MOI 1E-2, (**B**) MOI 1E-3, (**C**) MOI 1E-4, and (**D**) MOI 1E-5. The indicated signals correspond to FL and DI vRNAs. Upper thicker band of the ladder: 3000 bp, middle thicker band: 1000 bp, lower thicker band: 500 bp.

Taken together, our results demonstrate that the MOI has a strong effect on OP7 chimera DIP production. At high MOI, high fractions of OP7 chimera DIPs were present along with low total virus titers suggesting that OP7 chimera DIPs impeded virus propagation. On the other hand, higher total virus titers, but lower fractions of OP7 chimera DIPs were found at lower MOIs. Infections at intermediate MOIs of 1E-3 and 1E-4 appear to be a good compromise for achieving high OP7 chimera DIP fractions and high virus titers.

### The MOI used for production affects the *in vitro* interfering efficacy

Next, we aimed to identify the optimal MOI yielding OP7 chimera DIP material showing the highest *in vitro* interfering efficacy per product volume. Therefore, an *in vitro* interference assay (Fig. 4) was carried out to investigate the inhibition of STV propagation in the presence of OP7 chimera DIPs. In brief, WT MDCK(adh) cells were either infected with STVs only at a MOI of 10 (negative control, NC) or co-infected with 125 μL (fixed volume) of DIP material produced at different MOIs (Fig. 2).

**Figure 4:**
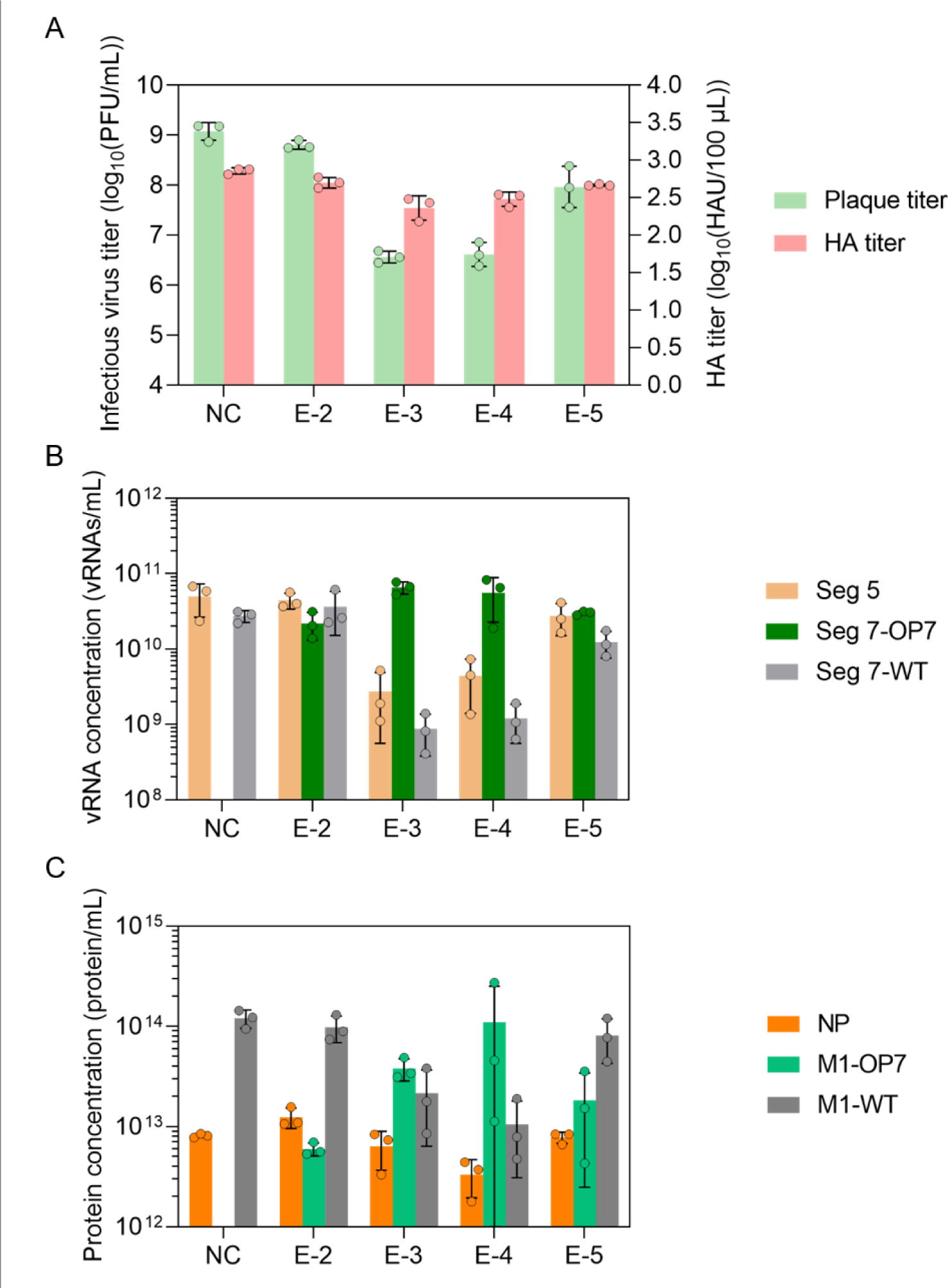
*In vitro* interference assay with OP7 chimera DIP material produced at different MOIs. MDCK(adh) cells were infected with STVs alone at a MOI of 10 (NC) or co-infected with 125 μL of indicated DIP material produced at different MOIs in shake flasks (Fig. 2). (**A**) Infectious virus release, indicated by plaque titer, and total virus release, indicated by HA titer, at 16 hpi. PFU, plaque-forming units; HAU, HA units. (**B**) Extracellular vRNA concentration, quantified by RT-qPCR. (**C**) Protein concentration, quantified by MS. NP, nucleoprotein. Interference assay was performed in three independent experiments, corresponding samples were quantified in a single measurement. Error bars indicate the standard deviation (SD). Samples of the optimal harvest time points (Fig. 2) were analyzed.

Our results indicated the strongest interfering efficacy for the OP7 chimera DIP material produced at a MOI of 1E-3 and 1E-4. This was shown by a suppression of the infectious virus release by more than two orders of magnitude (quantified by the plaque assay), which was significantly more than the decrease of only a factor of two, observed for the material produced at a MOI of 1E-2 (p < 0.0001, One-way analysis of variance (ANOVA) followed by Tukeýs multiple comparison test), and significantly different to the reduction of one log for the material produced at a MOI of 1E-5 (p < 0.001) (Fig. 4A). For the total virus release, as expressed by the HA titer (Fig. 4A) and extracellular Seg 5 vRNA concentration (Fig. 4B), this trend was less pronounced. Further, co-infections with highly interfering DIP material, produced at a MOI of 1E-3 and 1E-4, resulted in a pronounced OP7 phenotype, i.e. an overproportional extracellular Seg 7-OP7 vRNA concentration in comparison to other gene segments [19, 39], indicating the preferential replication of Seg 7-OP7 vRNA during virus propagation (Fig. 4B). Similarly, an overproportional M1-OP7 concentration relative to M1-WT was found in progeny virions (Fig. 4C).

In summary, the interfering efficacy of OP7 chimera DIP material strongly depended on the production MOI, with intermediate MOIs of 1E-3 and 1E-4 representing the optimum.

### High *in vivo* tolerability and antiviral efficacy of OP7 chimera DIP material

We next tested the tolerability and antiviral efficacy of produced OP7 chimera DIP in a mouse infection model. DIP material produced in shake flasks, purified by steric exclusion chromatography (SXC) [18, 19, 47, 48], was used (MOI 1E-4, 6.5 × 10^10^ virions/mL, c_DIP_ calculated based on HA titer, OP7 chimera DIP fraction of 60.07%, based on vRNA quantification). As a negative control, OP7 chimera DIP material that was inactivated with UV light for 24 mins (inactive, 5.4 × 10^10^ virions/mL) was utilized, which typically does not show an interfering efficacy *in vitro* [18, 19].

First, to test for the tolerability of the OP7 chimera DIP preparations, we administered 20 μL of active OP7 chimera DIPs (diluted to 1:2 and 1:20, corresponding to 6.5 × 10^8^ and 6.5 × 10^7^ virions per mouse, respectively) intranasally to the animals (Fig. 5A, B and C). Similar to PBS treatment, OP7 chimera DIP treatment was well-tolerated, as indicated by the absence of apparent weight loss (Fig. 5A) and clinical scores (Fig. 5B). Moreover, serum albumin levels in bronchoalveolar lavage (BAL) samples of PBS and OP7 chimera DIP-treated mice were comparable, indicating that OP7 chimera DIP administration did not compromise lung integrity (Fig. 5C). This was further confirmed by histopathological examination of the lungs. For mice treated with PBS (Fig. 6), only minimal interstitial pneumonia located near the hilus was observed; a finding that can be typically attributed to intranasal application of liquid to the lungs. Mice treated with OP7 chimera DIPs (1:2 and 1:20 dilution) showed a minimal increase in inflammatory infiltration. Yet, no histopathological changes that appear to be clinically relevant were observed, which is in line with the presentation of the clinical scores (Fig. 5B). Together, these data demonstrate that intranasal administration of OP7 chimera DIPs alone is well-tolerated.

**Figure 5:**
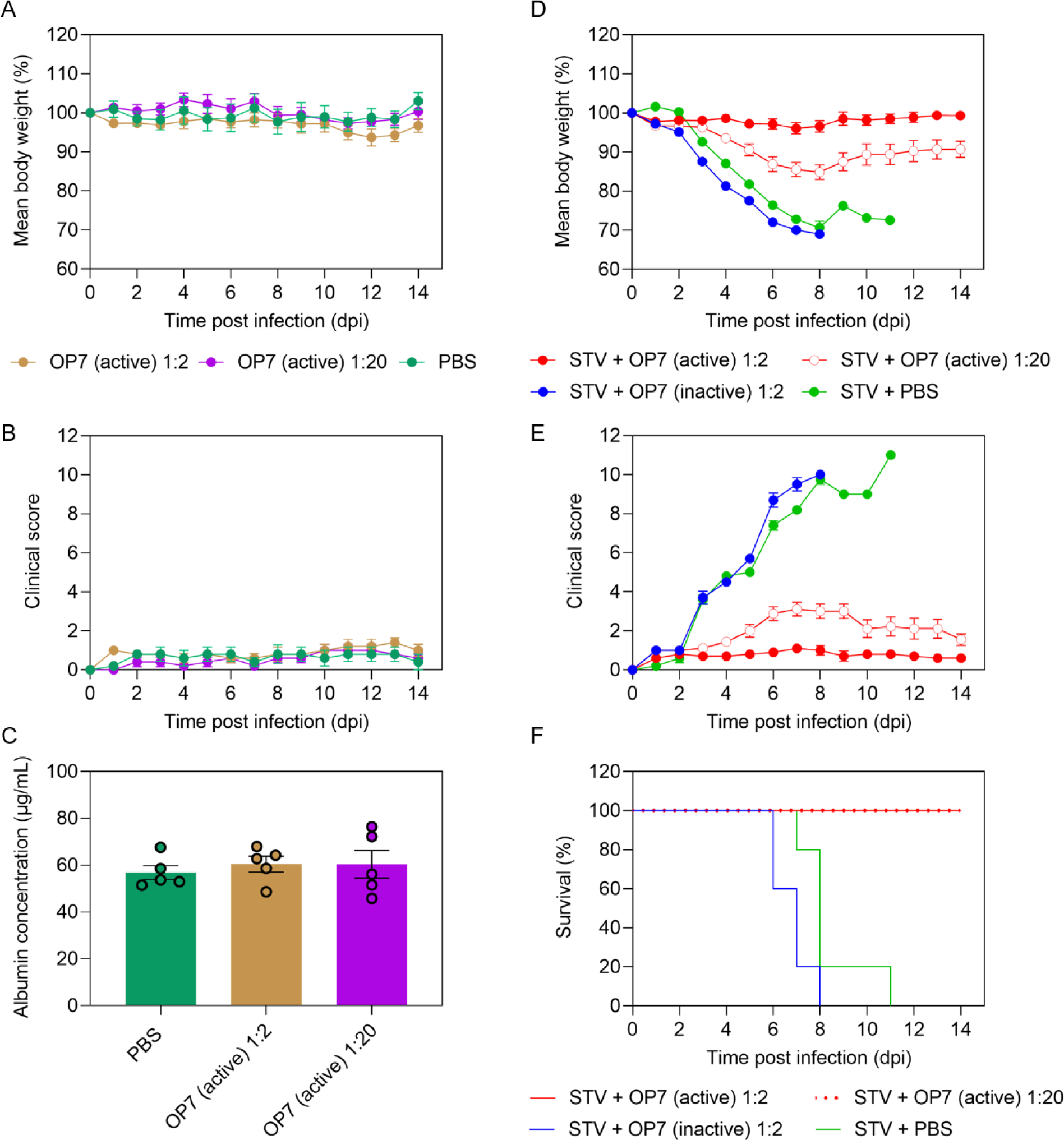
Antiviral efficacy of OP7 chimera DIPs in a mouse infection model. **(A**, **B** and **C**) 20 μL of active OP7 chimera DIP material diluted to 1:2 (6.5 × 10^8^ virions/mouse) or 1:20 (6.5 × 10^7^ virions/mouse), or 20 μL PBS was intranasally administered to 12–24 weeks old female D2(B6).A2G-Mx1^r/r^ mice (n=5). (**A**) Mean body weight loss. Two-way ANOVA and Tukey correction for multiple comparison did not reveal a significant difference between groups (p > 0.05). (**B**) Clinical score. (**C**) Serum albumin concentrations in BAL fluid were measured by ELISA at 14 dpi. One-way ANOVA did not reveal a significant difference between means (p > 0.05). (**D**, **E** and **F**) Mice were inoculated with a lethal dose of 1000 FFU of IAV STV (strain PR8) and co-treated with active OP7 chimera DIPs, diluted to 1:2 (n=10) or 1:20 (n=9), inactive OP7 chimera DIPs diluted to 1:2 (5.4 × 10^8^ virions/mouse, n=10) or PBS (n=5) in a total volume of 20 μL. (**D**) Mean body weight loss. The differences between the mean body weight of mice co-treated with 1:2 (mixed-effects model and Tukey correction for multiple comparison, p < 0.0001) or 1:20 OP7 chimera DIPs (p < 0.05) were significant relative to co-treatment with PBS. (**E**) Clinical score. (**F**) Kaplan-Meyer curve representing the survival rate. The differences between the survival of mice co-treated with 1:2 (log-rank test for two groups, p < 0.0001) or 1:20 OP7 chimera DIPs (p < 0.0001) were significant relative to co-treatment with PBS. (**F**) Clinical score. (**A-F**) Error bars indicate the standard error of the mean (SEM).

**Figure 6:**
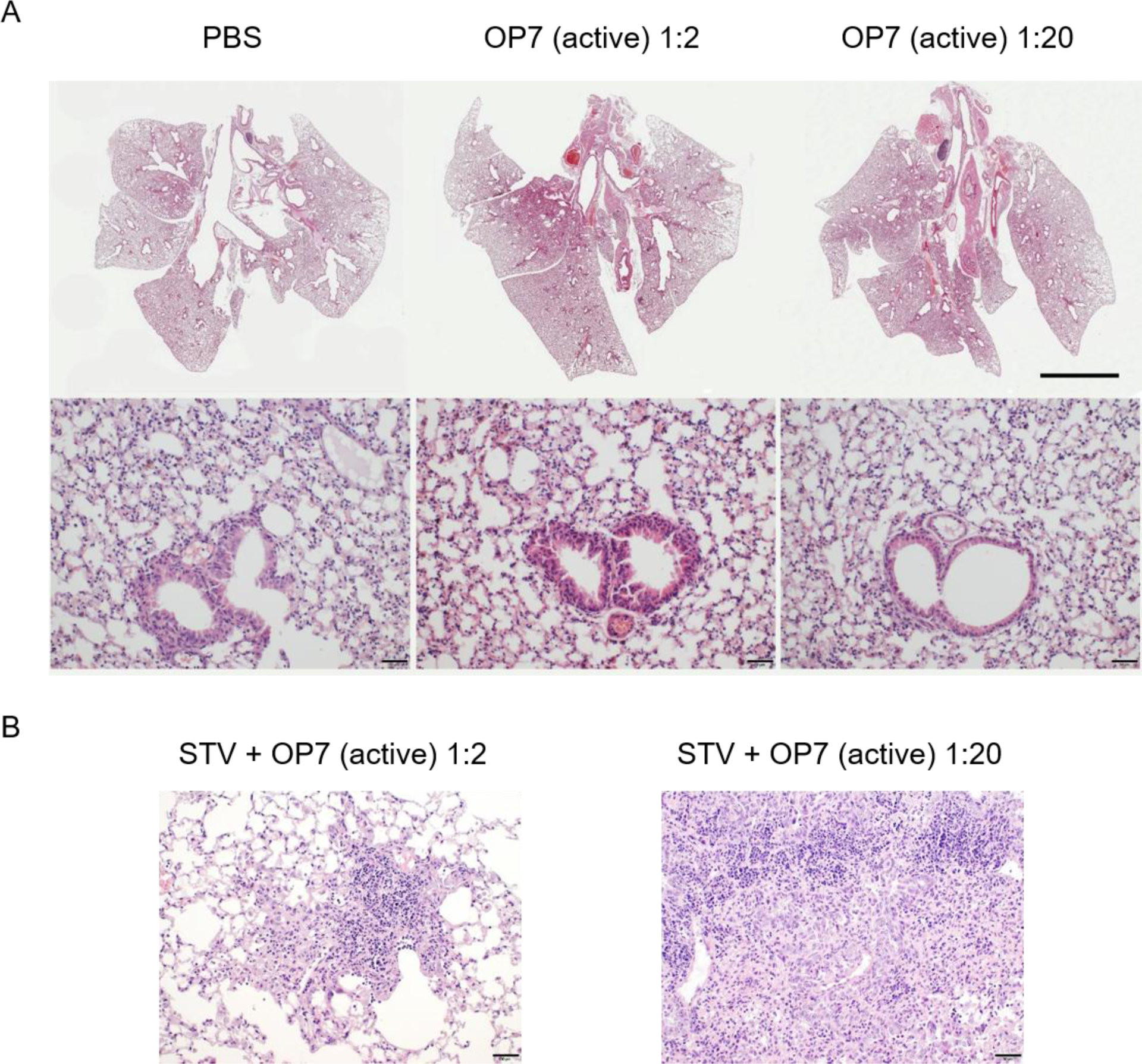
Histopathological changes in mouse lung sections after administration of OP7 chimera DIPs only or together with a lethal dose of STV. **(A**) 20 μL of active OP7 chimera DIP material diluted to 1:2 (6.5 × 10^8^ virions/mouse) or 1:20 (6.5 × 10^7^ virions/mouse), or 20 μL PBS was intranasally administered to 12–24 weeks old female D2(B6).A2G-Mx1^r/r^ mice (n=5). Histological images of the lungs after H&E staining in overview (top row, scale bar = 1 cm) and in detail (bottom row, scale bar = 50 µm). (**B**) Mice were inoculated with a lethal dose of 1000 FFU of STV of strain PR8 and co-treated with active OP7 chimera DIP material, diluted to 1:2 (6.5 × 10^8^ virions/mouse) or 1:20 (6.5 × 10^7^ virions/mouse) in a total volume of 20 μL. Histological images of the lungs after hematoxylin-eosin (H&E) staining in detail (scale bar = 50 µm).

Next, we infected mice with a lethal dose of 1000 focus-forming units (FFU) of IAV STV (strain PR8) and co-treated them with either active OP7 chimera DIPs (1:2 and 1:20), inactive OP7 chimera DIPs (1:2) or PBS (1:2) in a total volume of 20 μL. As expected, severe body weight loss and 100% IAV-induced mortality was observed in PBS co-treated mice (Fig. 5D and F). Further, similar to PBS co-treatment, mice co-treated with inactive OP7 chimera DIPs showed the same high infection-induced morbidity (Fig. 5D and E) and mortality (Fig. 5F), indicating the absence of protective efficacy by UV inactivated DIPs. In strong contrast, no body weight loss was observed when active OP7 chimera DIPs (1:2 diluted) were co-administered, while a higher dilution (1:20) resulted in a modest loss of body weight (approx. 16%). Importantly, all mice co-applied with active OP7 chimera DIPs (1:2 and 1:20) survived the otherwise lethal STV infection (Fig. 5F). Intriguingly, co-administration of active OP7 chimera DIPs (1:2) together with a lethal STV dose completely prevented the development of clinical signs (Fig 5E) related for influenza infection compared to PBS treatment only (Fig. 5B). Even co-administration of active OP7 chimera DIPs diluted to 1:20 together with a lethal dose of STV was highly effective in preventing a severe course of influenza disease (Fig. 5E). In line with these findings, histopathological analysis revealed only a low-grade pneumonia and pneumocyte type II hyperplasia of mice co-administered with a lethal dose of STVs and OP7 chimera DIPs at a dilution of 1:2 (Fig. 6). In comparison, a lethal dose of STV of strain PR8 typically resulted in hyper-inflammatory immune responses in infected lungs [49]. Even the co-application of the low (1:20) dose OP7 chimera DIPs was sufficient to protect the animals from a lethal outcome of pneumonia (Fig. 6).

Taken together, intranasal application of only the OP7 chimera DIP material is very well-tolerated in mice. Furthermore, co-administration of OP7 chimera DIPs mediated full protection against an otherwise lethal IAV STV infection. These data demonstrate the safety and a remarkable antiviral efficacy of the produced OP7 chimera DIPs *in vivo*.

## Discussion

Previous studies of our group showed that the MOI used for cell culture-based production of DIPs affects total virus yields [18, 19, 50, 51] and interference efficacy [18, 19]. Here, we found the highest interfering efficacy for material produced at intermediate MOIs of 1E-3 and 1E-4 (Fig. 2). For infection at the lower MOI of 1E-5, more single-hit infections might occur, in which propagation-incompetent OP7 chimera DIPs are not able to replicate. In addition, only low fractions of OP7 chimera DIPs and high total virus yields were observed in the harvest material. This suggests that Seg 1 cDIPs out-diluted OP7 chimera DIPs. On the contrary, at the higher MOI of 1E-2, more co-infections of Seg 1 cDIPs and OP7 chimera DIPs could occur due to higher virus concentrations. Accordingly, as a result of its superior replication, a higher fraction of OP7 chimera DIPs was observed (Fig. 2C and D). However, total virus yields were lower (Fig. 2B, C and D), likely due to the inhibiting effect of OP7 chimera DIPs on virus propagation. Therefore, at intermediate MOIs, a balanced trade-off between the fraction of OP7 chimera DIPs and total virus yields in the produced material appears to be decisive for the optimal interference efficacy observed.

Next, we evaluated the tolerability and efficacy of OP7 chimera DIP preparations harvested from shake flasks using mice. Most laboratory mouse strains lack the *Mx1* gene, which is an important IFN-induced restriction factor against IAV infections in mice and in humans. Therefore, we used a mouse model expressing a functional *Mx1* gene, called D2(B6).A2G-*Mx1^r/r^* [49]. Accordingly, this model better represents the immune response in humans to IAV infections. Intranasal administration of high doses of only OP7 chimera DIPs did neither result in disease, nor in clinically relevant histopathological changes in mice lungs, indicating a high tolerability after OP7 chimera DIP treatment (Fig. 6). These results (and those of other groups [5, 16, 20–25, 40]) clearly suggest that common concerns regarding adverse effects (e.g., cytokine storm, lung damage) due to DIP administration in animals can be abandoned and that DIPs, as defective, non-replicating viral particles might also be suitable for safe clinical applications in humans. Moreover, co-infection with a lethal dose of STV together with OP7 chimera DIPs resulted in 100% survival of the mice (Fig. 5F), and these animals did not even show signs of clinical disease (Fig. 5E). A similarly high antiviral activity was found for DIPs derived from IAV and from other viral species in different animal models [5, 16, 18–25, 40]. Together, these results demonstrate the safety and antiviral activity of DIPs and the potential for their *in vivo* use as antivirals.

In light of imminent pandemic threats, new broadly-acting antivirals that are readily available at low costs are required. IAV DIPs typically suppress a wide range of IAV strains including contemporary human epidemic, pandemic and even highly pathogenic avian IAV as demonstrated *in vitro* and in mouse and ferret experiments [16, 21, 25, 39, 40]. Surprisingly, IAV DIPs can even suppress unrelated virus replication. This unspecific protection is mediated by the ability of DIPs to stimulate innate immunity and to establish a so-called antiviral state. For instance, mice were rescued from a lethal dose of influenza B virus and pneumonia virus of mice by DIP co-administration [38, 41]. In addition, we demonstrated a pronounced antiviral effect against SARS-CoV-2 [37] and against respiratory syncytial, yellow fever and Zika virus replication *in vitro* (Pelz et al., submitted). Such broad protective immunity against many different unrelated viruses was also observed for dengue and poliovirus DIPs [24, 52]. This suggests that DIPs could be used as broadly-acting antiviral agents to treat viral infections as a fast countermeasure to protect people at risk and restrict virus spreading, e.g., in the case of a pandemic.

In our future studies, a scalable cell culture-based production and purification process for OP7 chimera DIPs should be established in bioreactors to achieve even higher titers and improve the purity of OP7 chimera DIP preparations. To leverage the antiviral potential of OP7 chimera DIPs, e.g. for use as an intranasal droplet spray [26, 27], the establishment of a good manufacturing practice (GMP) production process would then be the base for toxicology and safety studies, clinical trials and later on to market approval.

## Materials and Methods

### Cells and viruses

MDCK(adh) cells (ECACC, #84121903) and MDCK-PB2(adh) cells (expressing IAV PB2, generated by retroviral transduction, as described previously [43]) were maintained in Glasgow Minimum Essential Medium (GMEM, Thermo Fisher Scientific, #221000093) supplemented with 10% fetal bovine serum (FBS, Merck, #F7524) and 1% peptone (Thermo Fisher Scientific, #211709). Puromycin (Thermo Fisher Scientific, #A1113803) was added to a concentration of 1.5 µg/mL for MDCK-PB2(adh) cells. HEK-293T-PB2(adh) cells (expressing IAV PB2, generated previously [43]) were cultured in Dulbecco’s Modified Eagle Medium (DMEM) supplemented with 10% FBS, 1 % penicillin/streptomycin (10,000 units/mL penicillin and 10,000 µg/mL streptomycin, Thermo Fisher Scientific, #15140122) and puromycin at a concentration of 1 µg/mL. All adherent cells were maintained at 37 °C and 5% CO_2_.

MDCK-PB2(sus) cells (expressing IAV PB2, previously generated by retroviral transduction [18, 43]) were grown in chemically defined Xeno^TM^ medium (Shanghai BioEngine Sci-Tech), supplemented with 8 mM glutamine and 0.5 μg/mL puromycin. Cultivation of the suspension cells was performed in shake flasks (125 mL baffled Erlenmeyer flask with vent cap, Corning, #1356244) in 50 mL working volume in an orbital shaker (Multitron Pro, Infors HT; 50 mm shaking orbit) at 185 rpm, 37 °C and 5% CO_2_. To quantify VCC, viability and diameter of cell, Vi-cell^TM^ XR (Beckman Coulter, #731050) was used. IAV strain PR8 (provided by Robert Koch institute, #3138) was used for the interference assay. MOIs were based on the TCID_50_ titer for STV [53] (interference assay) or the plaque assay (OP7 chimera DIP production).

### Rescue of OP7 chimera DIP

The generation of OP7 chimera DIPs was based on a previously established plasmid-based reverse genetics system for the rescue of PR8-derived Seg 1 cDIPs [43]. Here, to complement the missing PB2 protein (deleted in Seg 1 cDIPs), a co-culture of HEK-293T-PB2(adh) cells and MDCK-PB2(adh) cells were used for plasmid transfections. For rescue of OP7 chimera DIPs, a pHW-based plasmid [54] harboring the sequence of Seg 7-OP7 (GenBank accession number: MH085234) was newly generated and kindly provided by Stefan Pöhlmann and Michael Winkler (German Primate Center, Goettingen, Germany). 50 ng of this plasmid was co-transfected with 500 ng of a pHW-based plasmid harboring the deleted Seg 1 sequence of a previously described cDIP (“Seg 1 gain” [44]) and 1 µg of the remaining plasmids for Seg 2-6 and 8 (pHW192-pHW196 and pHW198 [54], respectively) via the calcium phosphate-mediated transfection method. After reconstitution, OP7 chimera DIP material was amplified in MDCK-PB2(adh) cells and later used to infect MDCK-PB2(sus) cells for seed virus production.

### OP7 chimera DIP production in shake flasks

Production of OP7 chimera DIPs in shake flasks using MDCK-PB2(sus) cells was conducted with complete medium exchange prior to infection as described previously [18, 42]. In brief, cells in exponential growth phase were centrifuged (300 × g, 5 mins, room temperature) and resuspended in fresh medium (without puromycin) containing trypsin (final activity 20 U/mL, Thermo Fisher Scientific, #27250-018) at 2.0 × 10^6^ cells/mL. Cells were infected at different MOIs ranging from 1E-2 to 1E-5 at 37 °C and 5% CO_2_. At indicated time points, samples were centrifuged (3000 × g, 4 °C, 10 mins) and supernatants were stored at −80 °C until further analysis. RNA of progeny virions was extracted from supernatants using the NucleoSpin RNA virus kit (Macherey-Nagel, #740956) according to the manufacturer’s instructions and stored at −80 °C until PCR-based analysis.

OP7 chimera DIP material for mouse infection studies was produced in shake flasks at a MOI of 1E-4. Harvested DIP material was clarified (3000 × g, 10 mins and 4 °C) and sucrose (Merck, #84097) was added at a final concentration of 4 %. Next, the material was purified and concentrated by SXC as previously described [18, 19, 47]. Part of the purified, concentrated and sterile filtered DIP material was UV inactivated for 24 mins. Active (no UV inactivation), inactive (UV inactivated) DIP material and PBS spiked with sucrose (4% final concentration) were stored at −80 °C until further use.

### Virus quantification

Infectious virus titers were quantified using the plaque assay as previously described [18, 19, 39] using MDCK(adh) cells (interfering assay) or MDCK-PB2(adh) cells (determination of OP7 chimera DIP seed virus titer). Infectious virus titers were expressed as plaque-forming units (PFU)/mL. Furthermore, total virus concentrations were quantified using the HA assay as previously described [55]. Total DIP concentrations *C*_*DIP*_ (expressed as virions/mL) for the mouse infection studies were calculated based on the HA titer, and the concentration of the chicken red blood cells *C*_*RBC*_ used in the HA assay.

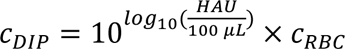

### Segment-specific RT-PCR

To detect contaminating DI vRNAs in Seg 2-Seg 8 in progeny virions, purified extracellular RNA was subjected to segment-specific PCR as described previously [39, 50]. In brief, RNA was reverse transcribed to cDNA using a universal “Uni12” primer [56] that binds to all eight genome segments. The resulting cDNA samples were used to amplify each genomic segment individually using segment-specific primers. PCR products were analyzed by agarose gel electrophoresis.

### RT-qPCR

In order to quantify the vRNAs purified from progeny virions, we used a previously described RT-qPCR method that enables polarity- and gene-specific quantification of individual vRNAs [19, 39, 50]. For this, a methodology involving tagged primers was employed [57]. Primers used for quantification of the vRNA of Seg 5 are listed in [39, 50], for Seg 7-OP7 in [19], and for Seg 7-WT, new primers were designed for the present study (for reverse transcription, Seg 7-WT tagRT for: 5’-ATTTAGGTGACACTATAGAAGCGTCTCGCTATTGCCGCAAA-3’ and for qPCR, Seg 7-WT realtime rev: 5’-CCTTTCAGTCCGTATTTAAAGC-3’). In order to allow for absolute quantification, RNA reference standards were used. vRNA concentrations were calculated based on calibration curves.

### Interference assay

The produced OP7 chimera DIP material was tested for the interfering efficacy *in vitro* according to a previously established protocol [19, 39]. Here, we assessed the inhibition of STV propagation upon co-infection with OP7 chimera DIP preparations. After infection, supernatants were analyzed for infectious and total virus titers using the plaque and HA assay, respectively. RT-qPCR and MS were used for quantification of vRNA and viral protein, respectively, of the progeny virions.

### Quantification of IAV proteins

MS analysis was used for absolute quantification of M1-WT, M1-OP7 and nucleoprotein (NP) according to a method described previously [45]. For this, we used isotopically labelled peptides of synthetic origin of corresponding proteins that were added as an internal standard before tryptic digestion of the samples for absolute quantification (AQUA). For M1-OP7, a peptide containing one mutation, which is not present in the M1-WT (EITFYGAK) was used. For quantification of M1-WT, two peptides exclusive for M1-WT (LEDVFAGK, QMVTTTNPLIR) were used. In brief, supernatant samples containing DIPs were heat inactivated (3 mins, 80°C) for further processing. Next, total protein concentration was determined using a Pierce^®^ BCA protein assay (Thermo Fisher Scientific, #23227,) according to the manufacturer’s protocol. Sample preparation for MS analysis was performed by using filter-aided sample preparation as described previously [45, 58]. After drying of the eluted peptides, 80 μL of mobile phase A (LC-MS-grade water, 0.1% trifluoroacetic acid) and 20 µL (=2 pmol of each peptide) of peptide standard mix containing isotopically labelled peptides of synthetic origin for M1-WT, M1-OP7 and NP were added to each sample. Subsequently, MS analysis was carried out as described before [45]. Raw files from Bruker timsTOF Pro were analysed by using Skyline (vs. 19.1) [59]. Absolute protein copy numbers and virus concentrations were calculated as described previously [45].

### Mouse infection experiments

D2(B6).A2G-Mx1*^r/r^* mice were generated by backcrossing DBA/2JRj mice for 10 generations onto congenic B6.A2G-Mx1r/r mice as described previously [49]. Mice were bred and maintained in individually ventilated cages in a specific pathogen-free environment (animal facility, Helmholtz Centre for Infection Research, Braunschweig Germany); food and water were provided *ad libitum*. Female, age-matched (12–24 weeks) D2(B6).A2G-Mx1*^r/r^* mice that harbor a functional MX dynamin-like GTPase 1 (Mx1) resistance gene were randomly allocated into experimental groups. Following intraperitoneal injection of ketamine/xylazine, mice were intranasally administered with 20 µL of active OP7 chimera DIPs or PBS at indicated concentrations to test for the tolerability. Moreover, antiviral efficacy was studied by inoculation with a lethal dose of 1000 FFU of IAV STV strain PR8 and co-treatment with active OP7 chimera DIP at indicated concentrations, inactive OP7 chimera DIPs or PBS in a total volume of 20 μL. Determination of the FFU titer was conducted as described elsewhere [60]. Following administration, health status (body weight, appearance of fur, posture, activity) of mice was monitored at least once per day. In case humane endpoint criteria were reached, animals were humanely euthanized and the infection was recorded as lethal. Bronchoalveolar lavage (BAL) samples were harvested as described in [61].Serum albumin concentrations in BAL fluids were measured by ELISA (Fortis Life Sciences, #E90-134).

### Histopathological analysis

Complete lungs of the mice were routinely fixed in 4% formalin and embedded in paraffin. Sections with 5 µm thickness were cut, dewaxed, and stained with hematoxylin-eosin (H&E). Histopathological evaluation was performed in a blinded manner by a veterinary pathologist certified by the European College of Veterinary Pathologists.

### Statistical analysis

All the statistical analysis and graph generation were performed using GraphPad Prism 9 software.

## Data availability

Data generated during this study can be requested from the corresponding co-author upon request.

## Acknowledgments

For the excellent technical assistance we thank Claudia Best, Nancy Wynserski and Karin Lammert. Xeno^TM^ medium used in the study were kindly supplied by Shanghai BioEngine Sci-Tech and Prof. Tan from the East China University of Science and Technology. We would like to thank Prof. Stefan Pöhlmann and Michael Winkler from German Primate Center, Goettingen, Germany for providing WT IAV segments and Seg 7-OP7 vRNA encoding plasmids.

## Author Contributions

Conceptualization, T.D., L.P., M.D.H., S.Y.K., and U.R.; Formal analysis, L.P., J.B.; Funding acquisition, A.G, K.S., D.B. (Dunja Bruder), U.R.; Investigation, T.D., L.P., J.B., J.K., O.K., P.M., M.B.; Project administration, T.D., L.P., S.Y.K.; Supervision, D.B. (Dirk Benndorf), A.G., Y.G., D.B. (Dunja Bruder), S.Y.K., U.R.; Visualization, T.D., L.P., O.K., S.Y.K.; Writing – original draft, T.D.; L.P.; Writing – review & editing, T.D., L.P., J.B., J.K., O.K., P.M., M.B., M.D.H, A.G., D.B. (Dirk Benndorf), Y.G., D.B. (Dunja Bruder), S.Y.K., U.R..

## Approval for animal experiments

All *in vivo* experiments were conducted after review and approval of the study protocol by institutional (HZI) and regional ethical bodies (Niedersaechsisches Landesamt fuer Verbraucherschutz und Lebensmittelsicherheit, 33.19-42502-04-18/2922).

